# Interfacial tension driven open droplet microfluidics

**DOI:** 10.1101/2021.07.29.454194

**Authors:** Jian Wei Khor, Ulri N. Lee, Jean Berthier, Erwin Berthier, Ashleigh B. Theberge

## Abstract

We developed an open channel droplet microfluidic system that autonomously generates droplets at low Ca (~10^-4^-10^-3^) by leveraging competing hydrostatic and capillary pressure. With only our open channel polytetrafluoroethylene (PTFE) device, pipettes, and commercially available carrier fluid, we produce hundreds of microliter droplets; tubing, electronics, or pumps are not required, making droplet technology feasible for research labs without external flow generators. Furthermore, we demonstrated conceptual applications that showcase the process of droplet generation, splitting, transport, incubation, mixing, and sorting in our system. Unlike conventional droplet microfluidics, the open nature of the device enables the use of physical tools such as tweezers and styli to directly access the system; with this, we developed a new method of droplet sorting and transfer that capitalizes on the Cheerios effect, the aggregation of buoyant objects along a liquid interface. Our platform offers enhanced usability, direct access to the droplet contents, easy manufacturability, compact footprint, and high customizability. This design is a first step in exploring the space of power-free open droplet microfluidic systems and provide design rules for similar channel designs.

## Introduction

We show for the first time droplet generation by an open microfluidic channel using passive forces aloneat low Ca (~10^-4^-10^-3^); we also demonstrate downstream manipulations that are uniquely enabled by the open nature of the system. This work has potential to open new avenues in droplet microfluidics, a field which has grown immensely in the past decade.(*1–11*) Droplet microfluidics is an attractive technology because it is high throughput, miniaturizes chemical and biological processes, reduces reagent waste, and enables the use of precious or expensive reagents. The microliter to picoliter droplets act as chambers to conduct biological or chemical analyses and have many important applications in DNA sequencing, directed evolution, materials chemistry, and chemical reactions.(*1–11*) Additionally, droplet manipulations such as sorting, mixing, and splitting are empowering for expanding the versatility and potential applications of droplet microfluidics.(*1, 2, 5–11*)

In our system, we autonomously generate droplets in an open channel without pumps or tubing by leveraging the inherent hydrostatic pressure difference between two immiscible fluids (the fluorinated carrier phase and aqueous phase), capillary pressure, and spontaneous capillary flow (SCF) — flow induced by capillary action.(*12–17*) In our autonomous droplet generating system, the top side of the channel is exposed to air, rather than being enclosed by a ceiling, which gives access to the channel where conventional droplet microfluidics with enclosed channels does not. Open channels are advantageous because a researcher can directly pipette into the channel or add/retrieve droplets, solid objects such as magnetic beads, or tissue samples as they wish. Previous work in open droplet microfluidics includes the analysis of different modes of immiscible fluid plug flow in an open channel with various solvents as carrier fluid.(*18*) Droplet manipulations (splitting, incubating, and merging) in an open channel were investigated where the droplets were pipetted by hand into the channel at specified locations prior to addition of the carrier fluid.(*19*) Additionally, liquid handling systems have been used with open systems to keep flow rates constant for investigations of shear on cells in a hanging droplet(*20*) and to generate droplets in open channels with syringe pumps.(*21*) Our open channel droplet generator presented here is distinct from this prior work in that it shows, for the first time, we demonstrate autonomous generation of droplets from an open capillary flow without the direct actuation of a pipette to generate each droplet. This feature is particularly complex as typical passive pumping or capillary flows do not produce the pressures or shear forces sufficient to generate traditional shear-based droplet generation. Furthermore, we demonstrate this droplet generation with much more traditional and functional biphasic liquids such as HFE-500 fluorinated oil, as opposed to the traditional organic solvents that have been more typical for open biphasic flows previously.(*18, 19*) The use of fluorinated oils is particularly important for compatibility with biological experiments.(*2, 5, 7, 8, 22, 23*) The importance of manipulations of droplets for cell biology was highlighted in Soitu *et al.*(*1, 24–27*) and Li *et al.*(*1, 28–31*) where droplets were formed under an immiscible phase by segmenting aqueous solution with a physical stylus and simple pipetting in an open system. While Soitu *et al.* also developed a method to generate droplets in an open system, our system achieves similar droplet size (100 nL) without the use of pumps and tubing.(*1, 24–27*) Other methods to generate droplets using pipettes include simple agitation of a biphasic solution either with or without cavity-containing microparticles to create an emulsion of droplets in an immiscible phase.(*1, 32, 33*) Additionally, there are methods of autonomous droplet generation utilizing a closed channel but they have limited downstream analysis and manipulation capabilities.(*34, 35*)

Herein we describe our autonomous droplet generation system that achieves 100 nL - 120 μL size droplets in an open microfluidic channel. We developed a theoretical model to determine the conditions for droplet formation which is derived from the difference between hydrostatic pressure and capillary pressure across the aqueous plug. Both the experimental results and theoretical model show that interfacial tension, contact angle, and constriction width play critical roles in droplet generation. Thus far our open microfluidic system has only been used to generate droplets containing water and culture media, and additional work is needed to optimize the system with droplets containing cells and other biological materials. In fact, the goal of future studies is to implement open droplet technology with biological samples. As a first step toward this goal we focused on the physical parameters needed for droplet generation and developed a suite of droplet manipulation techniques that capitalize on the open nature of our devices, including novel methods for moving droplets with a specialized stylus or tweezers to sort and transfer droplets within a device or into another device or well plate. Applications best-suited for our medium-throughput droplet generator will leverage the ability to easily manipulate hundreds to thousands of droplets, scaling droplet generation through parallelization without pumps if needed.

## Results

### Autonomous droplet generation in an open microfluidic channel

Our open droplet generation platform demonstrates the ability to autonomously generate droplets at 2 droplets/second without intervention, and the open surface nature of our channel provides direct access to manipulate droplets (Figure 1). There are five regions in the device; the origin of flow is at the (1) inlet reservoir which leads to the (2) converging region, followed by the (3) narrow constriction, (4) diverging region, and (5) outlet reservoir (Figure S1). The operation of the device is simple; the aqueous solution is pipetted into the converging region creating an aqueous plug. The anterior end of the plug meets the narrow constriction and the plug is in full contact with the channel walls (Figure 1d, left). Next, the carrier fluid, a fluorinated oil, is pipetted in the inlet reservoir to establish a hydrostatic pressure that pushes the aqueous plug into the narrow constriction (0.2 mm - 3 mm in width) (Figure 1d, middle). As the anterior portion of the aqueous plug exits the narrow constriction, it pinches off into small droplets in the diverging region and flows to the outlet reservoir (Figure 1d, right). The angles of the converging and diverging region (45° and 20° respectively) were found to generate the most consistent and monodisperse droplets. Furthermore, a pair of square protrusions forms the narrow constriction which creates a large capillary pressure to oppose the hydrostatic pressure of the carrier fluid at the inlet reservoir. Hydrostatic pressure is governed by the carrier fluid height (determined by the dimensions of the device) and density, while capillary pressure is governed by interfacial tension, contact angle, and meniscus radius of curvature. As carrier fluid fills the inlet reservoir the hydrostatic pressure increases. At the narrow constriction and converging-diverging region there is a 0.2 mm tall step with grooves on both sides to allow carrier fluid to flow past the aqueous plug and prewet the channel walls prior to droplet generation. The low contact angle of the carrier fluid on the polytetrafluoroethylene (PTFE) channel surface enables prewetting of the channel walls by spontaneous capillary flow (SCF) which was found to be essential for continuous droplet generation.

**Figure 1.**
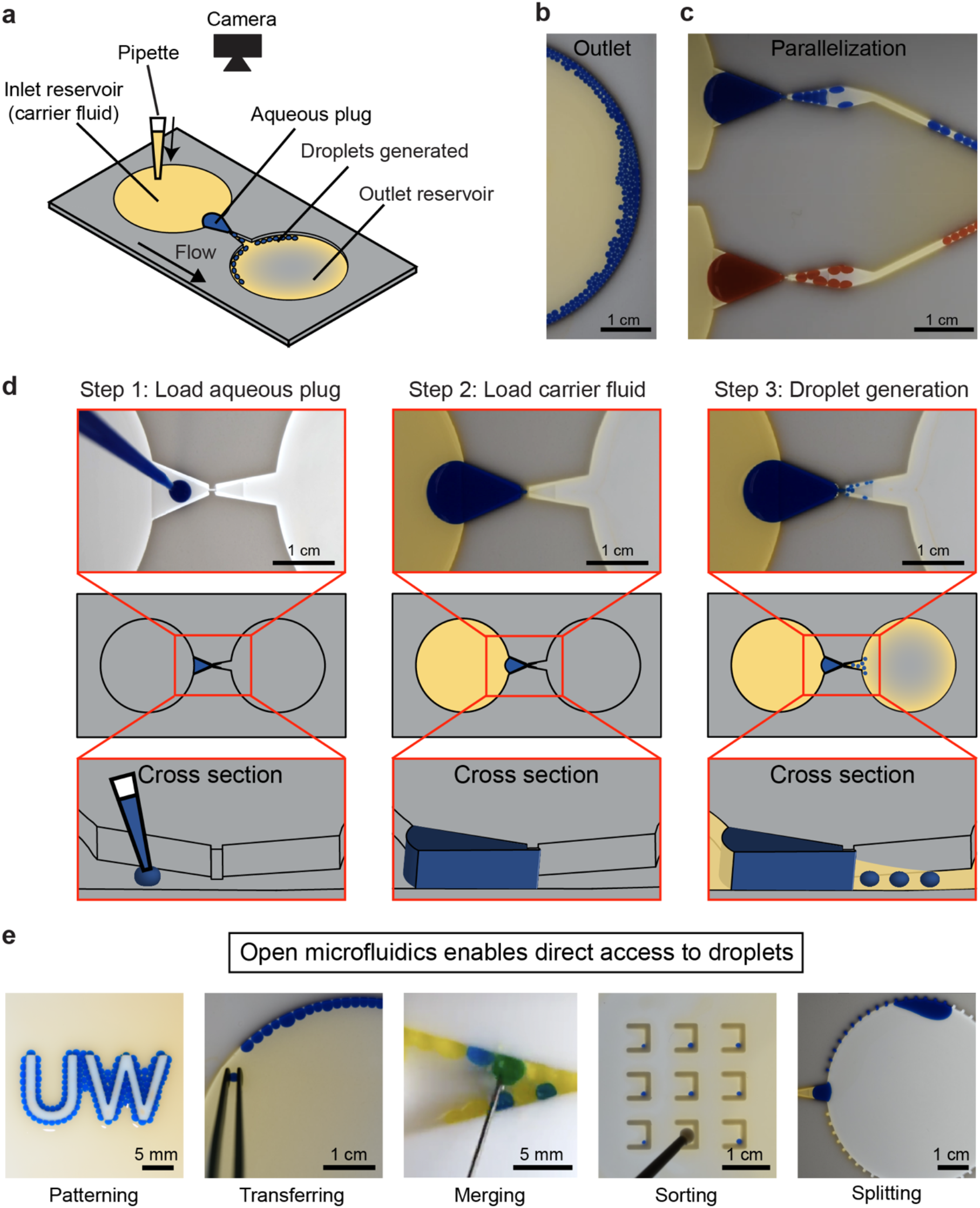
Device design, workflow, and manipulation of droplets. **a** Schematic representation of the device, **b** image of generated droplets in the outlet reservoir, and **c** droplets generated in parallel. **d** Workflow for droplet generation using passive forces derived from pressure (hydrostatic pressure and capillary pressure) in an open device (Supplementary Video 1). **e** Droplet manipulations downstream (Supplementary Video 2–6).

One critical breakthrough from our prior work is the use of PTFE as the channel material. Previously, in our open channel, channels were fabricated from poly(methyl methacrylate) (PMMA) and could only transport water droplets with organic solvents which is not ideal for applications of droplet microfluidics because organic solvents are often cytotoxic and known to remove small molecules from the droplets.(*18, 19*) In other words, using a carrier fluid that is not toxic and possess desirable interfacial properties was essential to furthering biological applications of our technology. The wetting properties of fluorinated oil, HFE 7500, on PTFE and the high contact angle of the aqueous plug allowed for droplet generation and manipulation (Figure 1) to occur in our channel. Fluorinated oils are ideal for life science applications in droplet-based microfluidics because of their biocompatibility—they have been used in applications ranging from human cell culture to digital droplet polymerase chain reaction (ddPCR).(*1–11, 22*)

### Condition for droplet generation

Droplet generation occurs when the aqueous plug overcomes the difference between hydrostatic and capillary pressure which pushes it into the narrow constriction. To note, a non-wetting plug should move spontaneously away from convergent geometries when no external forces of parallel direction are applied, and in our channel design, the hydrostatic pressure resulting from the carrier fluid is able to counteract this tendency.(*13*) Based on empirical observation from high-speed videos, the anterior portion of the aqueous plug is initially settled at the narrow constriction. As the anterior portion of the aqueous plug advances into the narrow constriction (plug advancement phase) it forms a bulbous shape (bulbing phase). Next, a liquid thread is formed (thread formation phase in Figure 2a and red arrow depicting the thread in Figure 2b), and the liquid thread diameter decreases until it pinches off and droplet formation occurs (pinch off phase).

**Figure 2.**
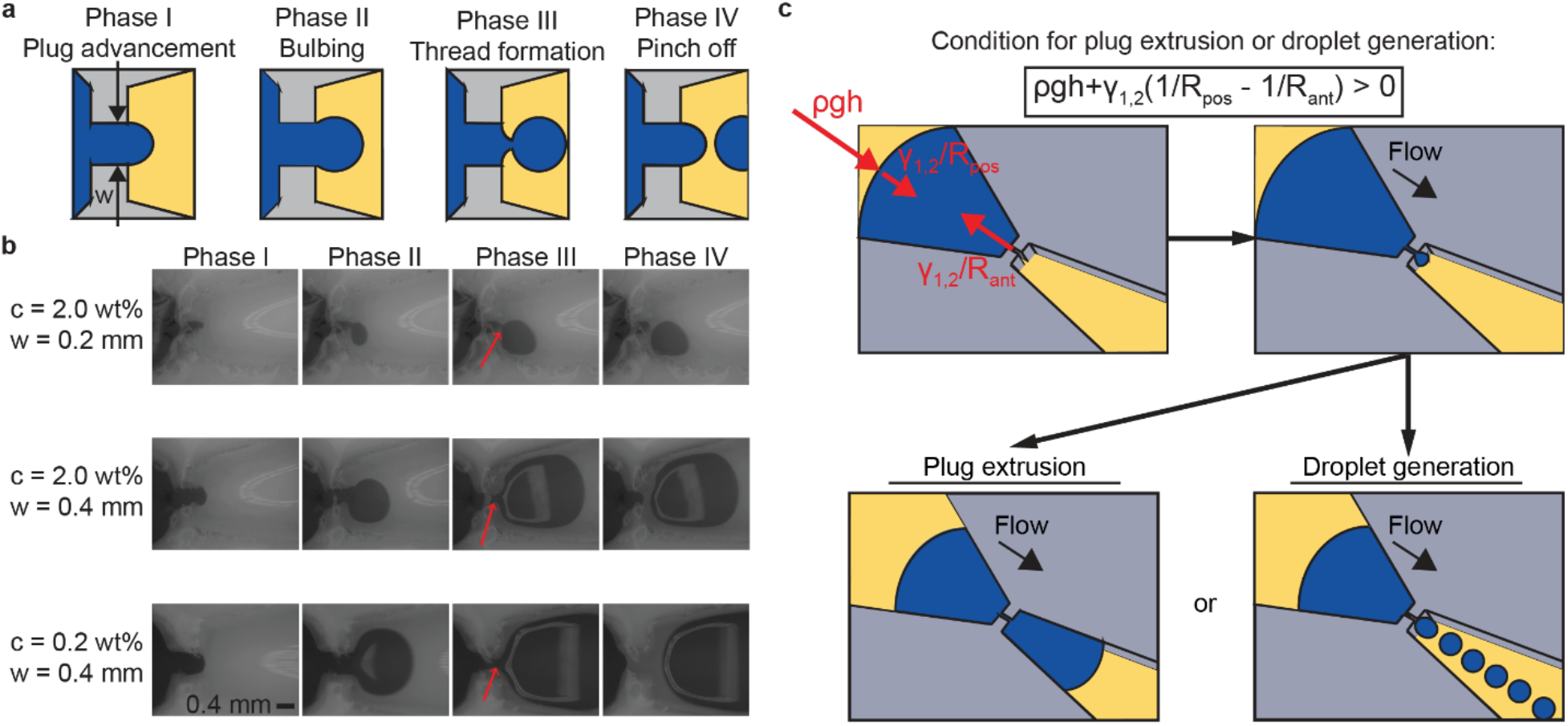
Snapshots of droplet formation with a high-speed camera and geometry of the open channel constriction for droplet generation. **a** Schematic of the phases observed during droplet formation. **b** Droplet formation in the constriction at various constriction width and surfactant concentration. Red arrow depicts the thread formation. **c** Hydrostatic pressure overcoming capillary pressure resulting in the aqueous plug being extruded or generating droplets at the constriction.

Figure 2b shows the droplet formation process for various surfactant concentrations, c, and constriction widths, *w*. We observed that the phases of the droplet formation process are nearly identical across the various surfactant concentrations and constriction widths used in this study (Figure 2b). We also observe that the constriction geometry and surfactant concentration is a determining factor in the final droplet volume generated. For the aqueous plug interface to extend past the narrow constriction, the hydrostatic pressure, *ρgh* at the posterior side of the plug must overcome the capillary pressure, *γ*_1,2_(1/*R*_*pos*_ − 1/*R*_*ant*_) (Figure 2c). Then, two possible outcomes occur: plug extrusion or droplet generation. We define plug extrusion as the entire aqueous plug passing through the constriction and sometimes breaking into two or three segments when exiting the diverging region. Droplet generation is a consistent generation of droplets pinching off from the aqueous plug with a relative standard deviation in volume of 5% - 7%. The pressure difference when the aqueous plug interface extends past the narrow constriction is:

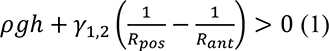

where *g* is the gravitational acceleration, *γ*_1,2_ is the interfacial tension between carrier fluid and aqueous plug, *ρ* is the carrier fluid density, *R*_*pos*_ is the radius of curvature of the posterior plug interface, and *R*_*ant*_ is the radius of curvature of the anterior plug interface. The left-hand side of Eq. (1) must be greater than 0 for the aqueous plug interface to extend past the narrow constriction.

By rearranging the pressure difference involving the hydrostatic pressures and capillary pressures, the condition for the aqueous plug extrusion or droplet generation is:

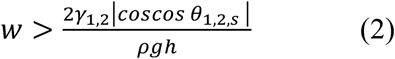

where w is the constriction width, *θ*_1,2,*s*_ is the contact angle between aqueous plug, carrier fluid, and channel wall, and *h* is the carrier fluid height. Based on experimental observation, the aqueous plug height is equal to the carrier fluid height. For the theoretical derivation, please see SI Supplementary Note 1.

We display the theoretical threshold for when the aqueous plug extrusion or droplet generation occurs in a regime map (Figure 3a dashed line, from Eq. (2)) at varying interfacial tension, contact angle, constriction width, and constriction height. In Figure 3a, we observed empirically that there exists a well-defined region (teal shaded region) where droplet generation occurs when constriction width is small (*w* ≤ 0.6 mm) and the condition in Eq. (2) is met (discussion on units for Eq. (2) can be found in SI Supplementary Note 2 and Figure S2). In the droplet generation region (teal shaded region), the constriction width *w*, the channel height *h*, interfacial tension *γ*_1,2_, and contact angle *θ*_1,2,*s*_ are the four factors that determine the volume of the droplets formed when carrier fluid density remains constant. The *γ*_1,2_ and *θ*_1,2,*s*_ is varied by the concentration of the surfactant, c. We observed that at a constriction width of 0.2 mm, when *γ*_1,2_ and *θ*_1,2,*s*_ increases (due to lowering fluorinated surfactant (FS) concentration from 2.0 wt% to 0.2 wt% or lower), the anterior portion of the aqueous plug cannot enter the narrow constriction and no plug extrusion occurs (Figure 3b). However, the aqueous plug can extrude from the narrow constriction if the constriction width increases to 0.4 mm at a FS concentration, c, of 0.2 wt%. This observation matches Eq. (2) and the regime map at Figure 3a; when *γ*_1,2_ and *θ*_1,2,*s*_ increases, the minimum width that allows plug extrusion is larger as well. Note that plug extrusion is possible without droplet generation (*w* = 3.0 mm, c ≥ 0.2 wt%). At the highest *γ*_1,2_ and *θ*_1,2,*s*_ (0.0 wt% FS), the anterior portion of the aqueous plug cannot enter the narrow constriction for any of the constriction width dimensions tested, thus plug extrusion does not occur (Figure 3b, consistent with the phase diagram in Figure 3a).

**Figure 3.**
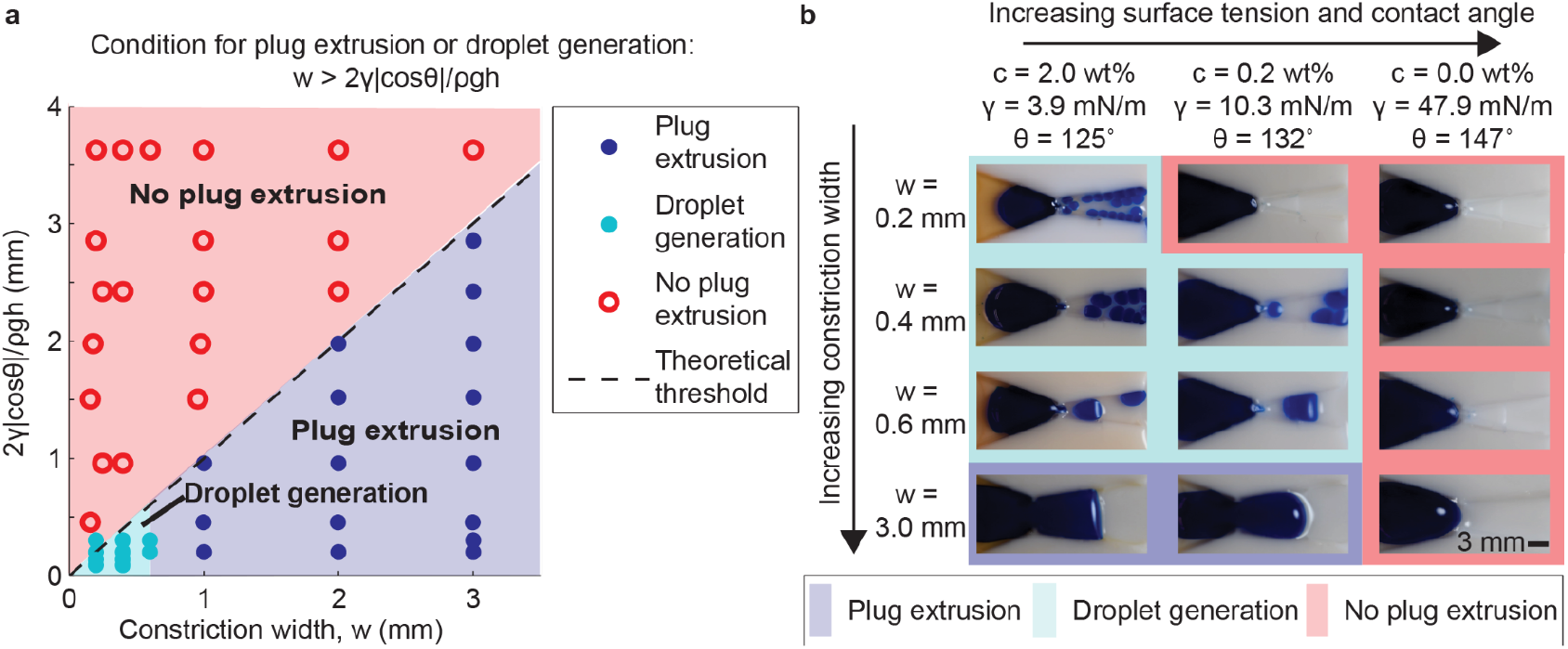
Conditions for plug extrusion and droplet generation. **a** Regime map of plug extrusion and droplet generation with theoretical threshold (dashed line per Eq. (2)); experimental data points of plug extrusion (purple filled circle), droplet generation (teal filled circle), and no plug extrusion (red open circle). The regions, predicted by the theoretical model, are plug extrusion (purple shaded region), droplet generation (teal shaded region), and no plug extrusion (red shaded region). Full table of parameters available in Table S1. **b** Montage of plug extrusion and droplet generation when varying constriction width, fluorinated surfactant concentration (c, wt%), interfacial tension *γ*_1,2_, and contact angle *θ*_1,2,*s*_.

Figure 4 shows the generated droplet volume as a function of constriction width, surfactant concentration, constriction length, and constriction height (data point presented as mean ± SD, n ≥ 90). Figure 4a depicts the effect of surfactant concentration (c = 2 wt% and c = 0.2 wt%) and constriction width (*w* = 0.2 mm and *w* = 0.4 mm) on generated droplet volume. The droplet volume generated increases with increasing constriction width *w* and decreasing surfactant concentration (interfacial tension *γ*_1,2_ and contact angle *θ*_1,2,*s*_).

**Figure 4.**
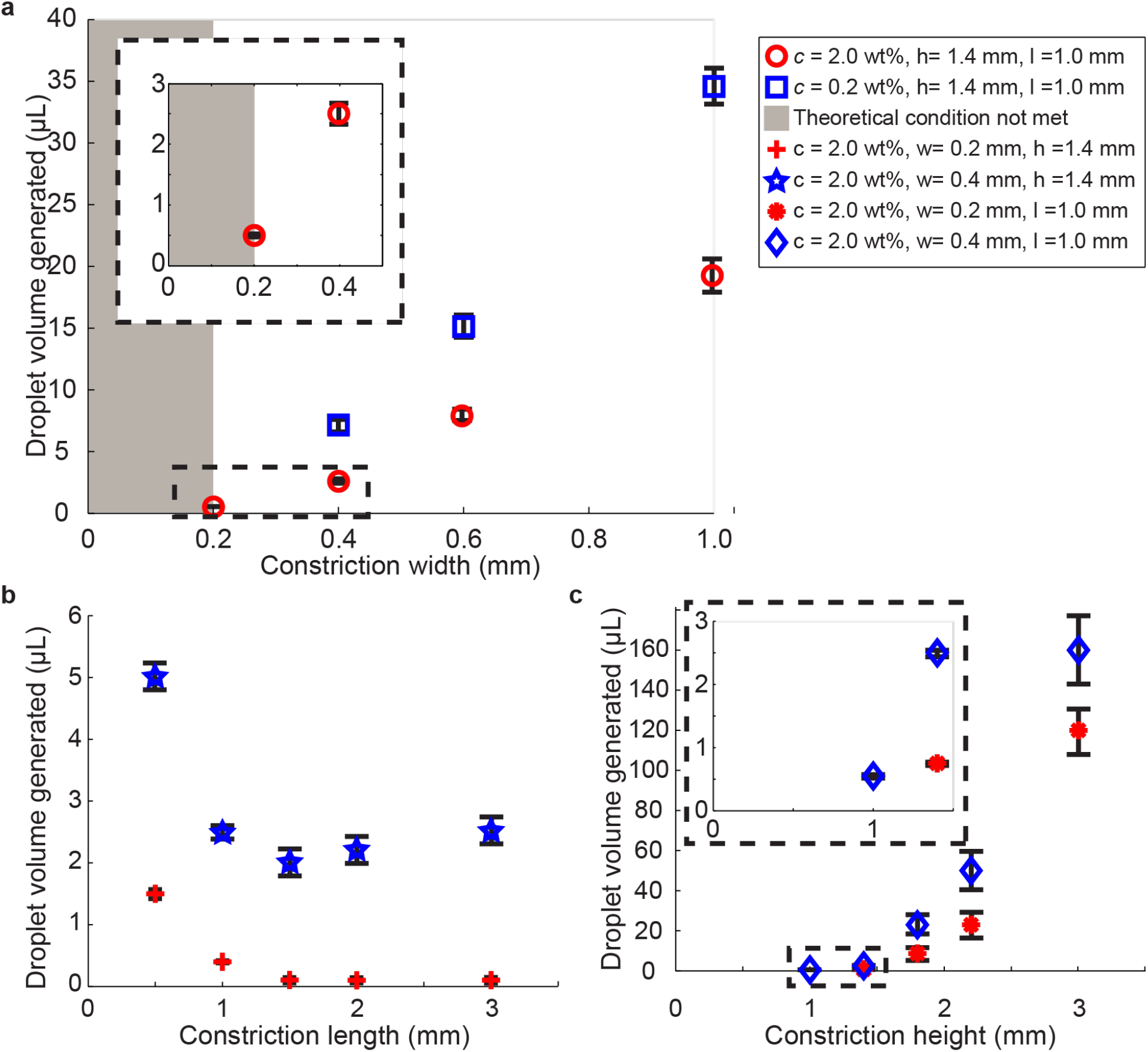
Plot of generated droplet volume as a function of **a** constriction width and surfactant concentration, **b** constriction length, and **c** constriction height. The shaded gray region is where the condition for the aqueous plug extrusion or droplet generation in equation (2) is not met. (Data point presented as mean ± SD, n ≥ 90). Table of parameters available in Table S2.

Similarly, when varying constriction length, *l* at *w* = 0.2 mm and *w* = 0.4 mm, we found that the generated droplet volume decreases as constriction length increases (Figure 4b). However, it is important to note that the standard deviation of the generated droplet volume is much larger when the constriction length is larger than 1 mm. When varying constriction height, *h* at *w* = 0.2 mm and *w* = 0.4 mm, we found that the generated droplet volume increases as constriction height increases (Figure 4b).

Overall, the smallest droplet volume generated in our experiments is 0.52 ± 0.025 μL and *w* = 0.4 mm is 2.39 ± 0.175 μL. The relative standard deviation of our droplet generation method ranges between 5% and 7% while conventional methods range between 1% and 3%(*36, 37*) which shows that the monodispersity of our system is relatively close to conventional methods, with room for improvement. We believe the relative standard deviation can be reduced if we have better control of maintaining relatively constant hydrostatic pressure, a focus of future redesigns. We also note that the physics underlying the droplet formation process is scalable to smaller length scales, therefore smaller droplet volumes will be achievable in future work if alternative fabrication processes such as advanced CNC milling or cleanroom fabrication processes are used to create devices with smaller constriction widths. Here, we chose 0.2 mm as the minimum constriction width due to the limitations of our current CNC milling process.

### Downstream droplet manipulation in open microfluidic devices

Droplet manipulation is essential for utilizing droplet microfluidics to study biology and chemistry. We demonstrate droplet manipulations to present the possibilities of conducting future biological and chemical studies with this device. In open systems, contamination and evaporation may be problematic if not controlled. Methods to address evaporation and contamination such as secondary containment and sacrificial liquid droplets(*38*) or wells have been successfully employed with primary human samples.(*39*) The open surface of the device enables us to create new methods to physically move the droplets such as using specialized tweezers or a needle to reach into the device and move or mix targeted droplets chosen in real-time at the discretion of the researcher. We use these manipulations for chemical reactions followed by droplet sorting.

Droplet transport and droplet sorting is an integral component of droplet microfluidics workflows. We developed a new method to selectively transfer single droplets to locations both on and off the open droplet generating device by picking them up with PTFE-coated tweezers or moving them with a PTFE stylus. The PTFE-coated tweezers and styli transport droplets laterally by utilizing the Cheerios effect, which is the phenomenon of multiple objects aggregating due to the objects deforming the liquid interface (Figure 5a).(*40–44*) The PTFE-coated tweezers and styli are placed at the surface of the carrier fluid, causing the carrier fluid to wet the tweezers or stylus and form a rising interface. The droplets are buoyant and when they are near the meniscus created by the tweezers or stylus they then move toward the upward rising fluid interface. Additionally, once the droplet is captured between the tweezers, the droplet rises up the tweezers due to the tendency for droplets to move towards divergent geometries (Figure 5b).(*13*) This allows droplets to be picked up and transferred to another location (e.g., well plate). To detach the droplet from the tweezers or stylus the user can abruptly move it away from the droplet which causes detachment due to inertia. Further, walls or other features (Figures 5c, 6b, and 6c) have menisci which enables droplets to again move up the rising interface to rest on the wall.

**Figure 5.**
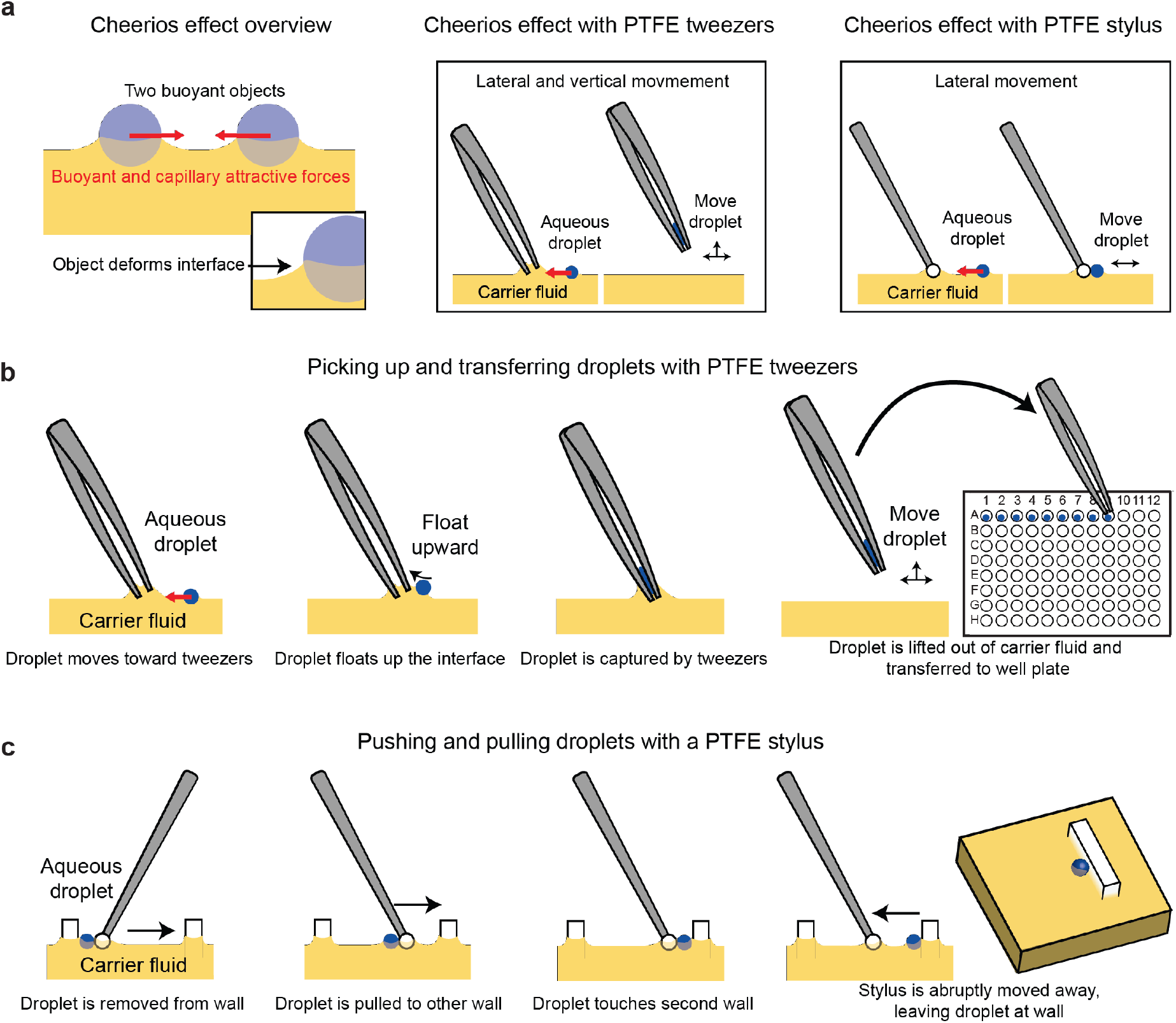
The Cheerios effect enables droplet transport. **a** Two neighboring buoyant objects aggregate due to the Cheerios effect.(*42, 44*) The buoyant object deforms the surface of the liquid resulting in a rising interface, and when a second buoyant object is near it tends to float up the rising interface. The tweezers and PTFE stylus behave similarly because they also deform the surface so the droplets tend to aggregate towards them. **b** Picking up droplets from an open microfluidic device and transferring them to another location (e.g., well plate) with PTFE-coated tweezers. **c** Transporting droplets from one wall to another with a PTFE ball mounted on a stylus.

PTFE tools are important for droplet transport because the PTFE prevents the aqueous droplet from strongly adhering to the tweezers, facilitating droplet release. Using a similar mechanism, we use a stylus with a PTFE bead adhered to the tip to gently move droplets around the reservoir for sorting and patterning droplets (Figure 6b and 6c). To summarize, the PTFE stylus enables lateral droplet transport and the PTFE-coated tweezers enables both lateral droplet transport and vertical droplet transport (i.e., picking up droplets). In future work, this approach could be used to sort droplets based on a reaction outcome or transfer droplets to a different part of a chip for further use in analysis or reactions.

**Figure 6.**
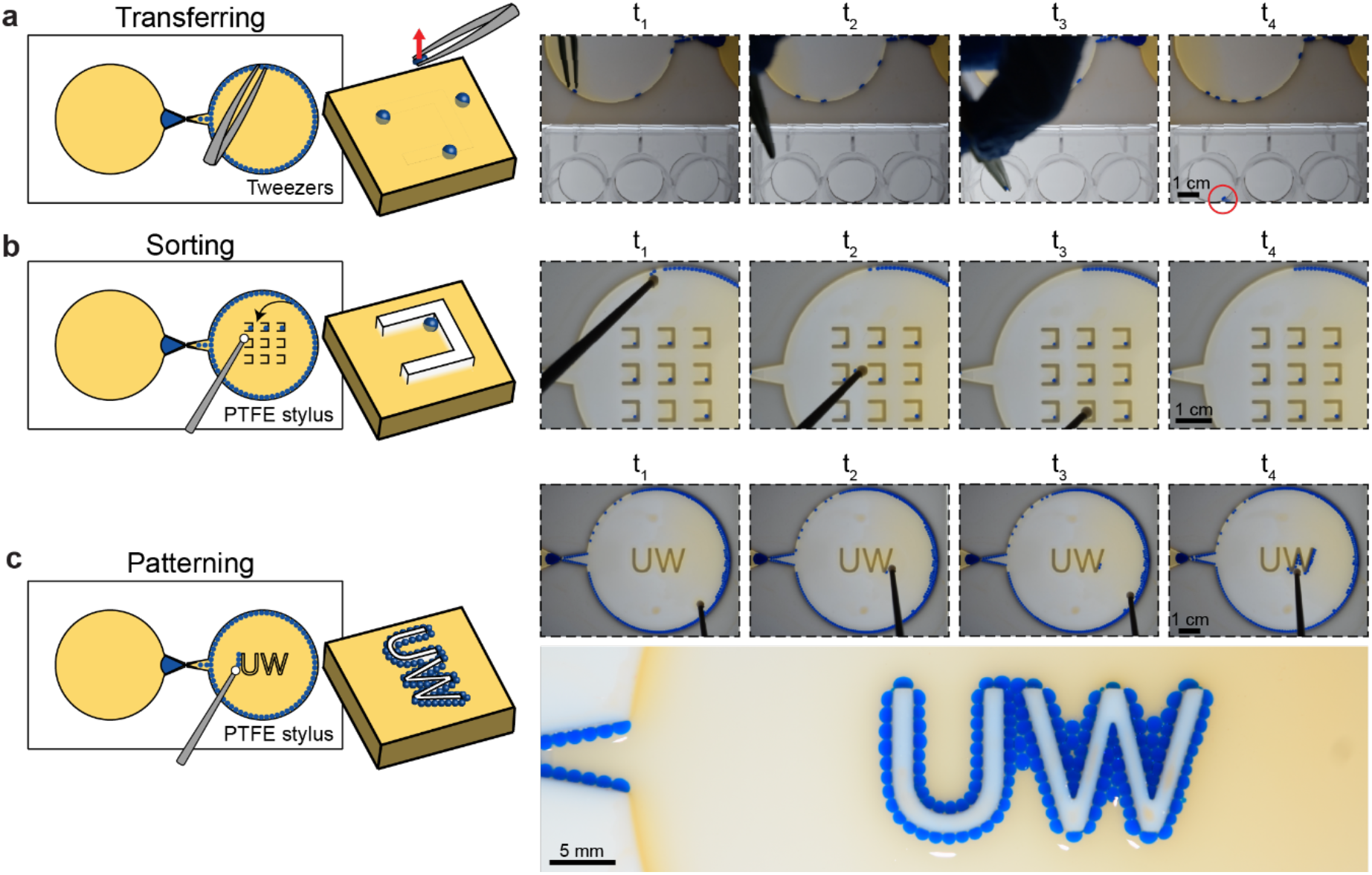
Open surface of the open microfluidic channel enables on-demand direct manipulation of droplets. **a** Selectively retrieving droplets from an open microfluidic device and transferring them to a well plate with PTFE-coated tweezers (Supplementary Video 3). **b** Sorting droplets into individual chambers with a PTFE ball mounted on a stylus (Supplementary Video 5). **c** PTFE stylus is used to transfer droplets to the “UW” pillar (Supplementary Video 2).

While we focused on moving individual droplets to highlight the ability to move droplets selectivity, this method can be multiplexed using an array of styli (for example, a three-pronged stylus could be used to move three droplets at once from the columns of the array similar to Figure 6c). The direct access to the droplets is unique to our open channel systems and allows users to directly extract and/or manipulate the contents of the channel. This is enabling for users who cannot rely on power-heavy solutions in conventional droplet microfluidics like acoustophoresis, electrophoresis, and electrowetting-on-dielectric (EWOD) or users who would like the flexibility to manipulate droplets at any point in their device, without the need to pre-pattern electrodes.(*10*)

For applications requiring automation, this method could be automated by mounting the stylus on a robotic XYZ controller interfaced with a fluorescent or colorimetric readout in the droplets to facilitate sorting based on a chemical/biochemical readout. The flexibility to move droplets on demand—both on- and off-chip—shows great promise for being able to conduct multistep chemical and biological experiments and incorporating analytical readouts that are best implemented outside of the original microfluidic chip embodiment.

In Figure 6c, the generated droplets are patterned by transporting them toward the “UW” feature using a PTFE bead to outline the feature. It is important to note that droplets were transported to sharp corners of the “W” with ease using the PTFE bead. In contrast, traditional droplet microfluidics would require pumps and valves to change the trajectory of the droplets. Furthermore, our channel design is not constrained by requiring prior knowledge of the location to which the droplets will be transported. This allows our channel design to be modular by being able to continuously upgrade or add new features (e.g., replacing “UW” feature with another feature) directly on the channel. Open microfluidics can be leveraged to make droplet studies user-friendly, customizable, and adaptable for integration of physical probes and tools downstream.

The ability to take small aliquots of a droplet after it is generated is useful for downstream analysis or processing steps, particularly when working with small volumes of precious reagents.(*1–11*) We demonstrated that we can split smaller droplets off from a large aqueous plug with a crenulation feature along the outlet reservoir wall (Figure 7a). In this case, the constriction allows aqueous plug to extrude and flow along the crenulations. As the plug travels along the crenulations via SCF, it fills the crenulation; as the posterior end of the large plug fully passes the crenulation it shears off leaving the crenulation filled with a small droplet. The droplets in the crenulations can then be retrieved with a pipette.

**Figure 7.**
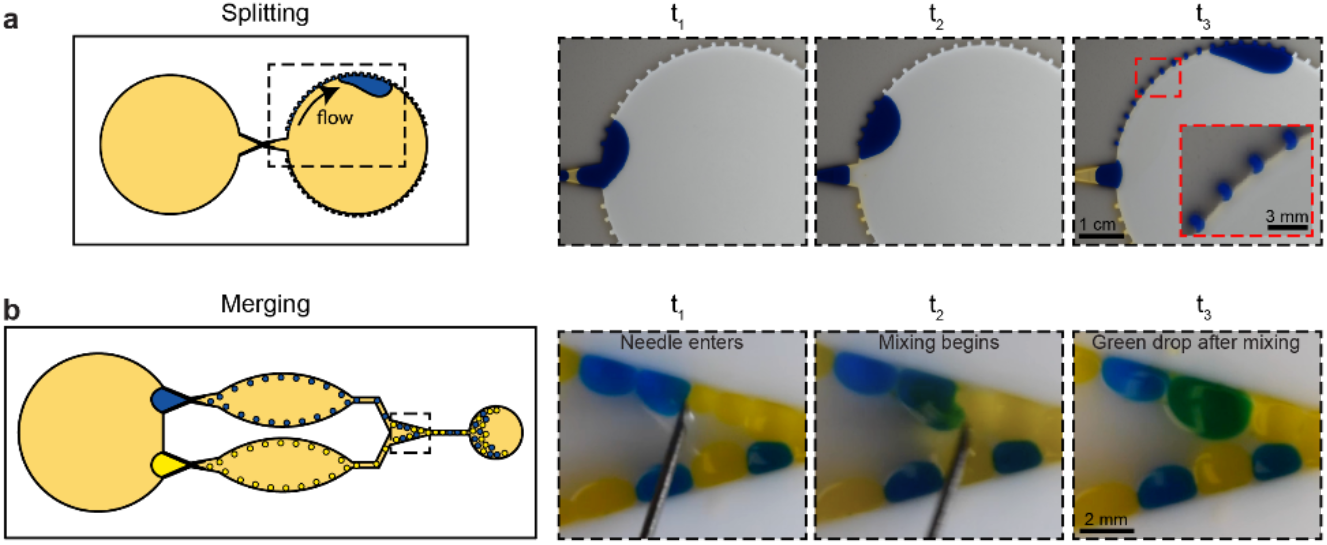
Splitting and merging droplets in an open droplet generator. **a** Crenulations along the outlet reservoir capture portions of an aqueous plug, and as the plug passes the crenulation, a small droplet splits off and remains in the crenulation (Supplementary Video 6). **b** A yellow and blue droplet are merged by mixing with a needle to form a green droplet (Supplementary Video 4).

In Figure 7b, two droplet merging is demonstrated by disrupting the interface of the droplets using a simple needle. The needle is moved back and forth between the two droplets to disrupt their interface resulting in their merging. In future work, the merging step could also be automated using a needle mounted on an XYZ controller as described for the droplet transport with tweezers and styli mentioned above. This merging technique with the automated system makes it comparable to prior droplet fusion methods (such as electrofusion or fusion based on electrowetting on dielectric), which often require the placement of specific components (e.g., electrodes) in the device design.(*1, 10*) Our new fusion method allows users to induce fusion on demand at any location in the open device; the location need not be decided prior to the experiment.

### Sorting reacted products in droplets

Droplet merging and sorting opens up applications for chemical reactions, biochemical assays, and other multi-reagent processes.(*45*) Selective droplet merging is demonstrated in Figure 8 using droplets of KSCN and Fe(NO_3_)_3_ mixed with blue dye for visualization. Droplets of the two reagents are formed using two parallel open-channel droplet generators. Then, two droplets are randomly selected from the outlet reservoir and transported to a platform with a pair of PTFE-coated tweezers (yielding three possible combinations: two droplets containing KSCN, two droplets containing Fe(NO_3_)_3_, or a droplet containing KSCN and a droplet containing Fe(NO_3_)_3_). On the platform, the droplets are merged by disrupting the interfaces of the droplet pair with a needle. When a droplet containing KSCN and a droplet containing Fe(NO_3_)_3_ merge, they change color from blue to red due to the formation of Fe(SCN)_3_. Merged droplets are then sorted to chambers categorized by reacted and unreacted droplets using PTFE-coated tweezers.

**Figure 8.**
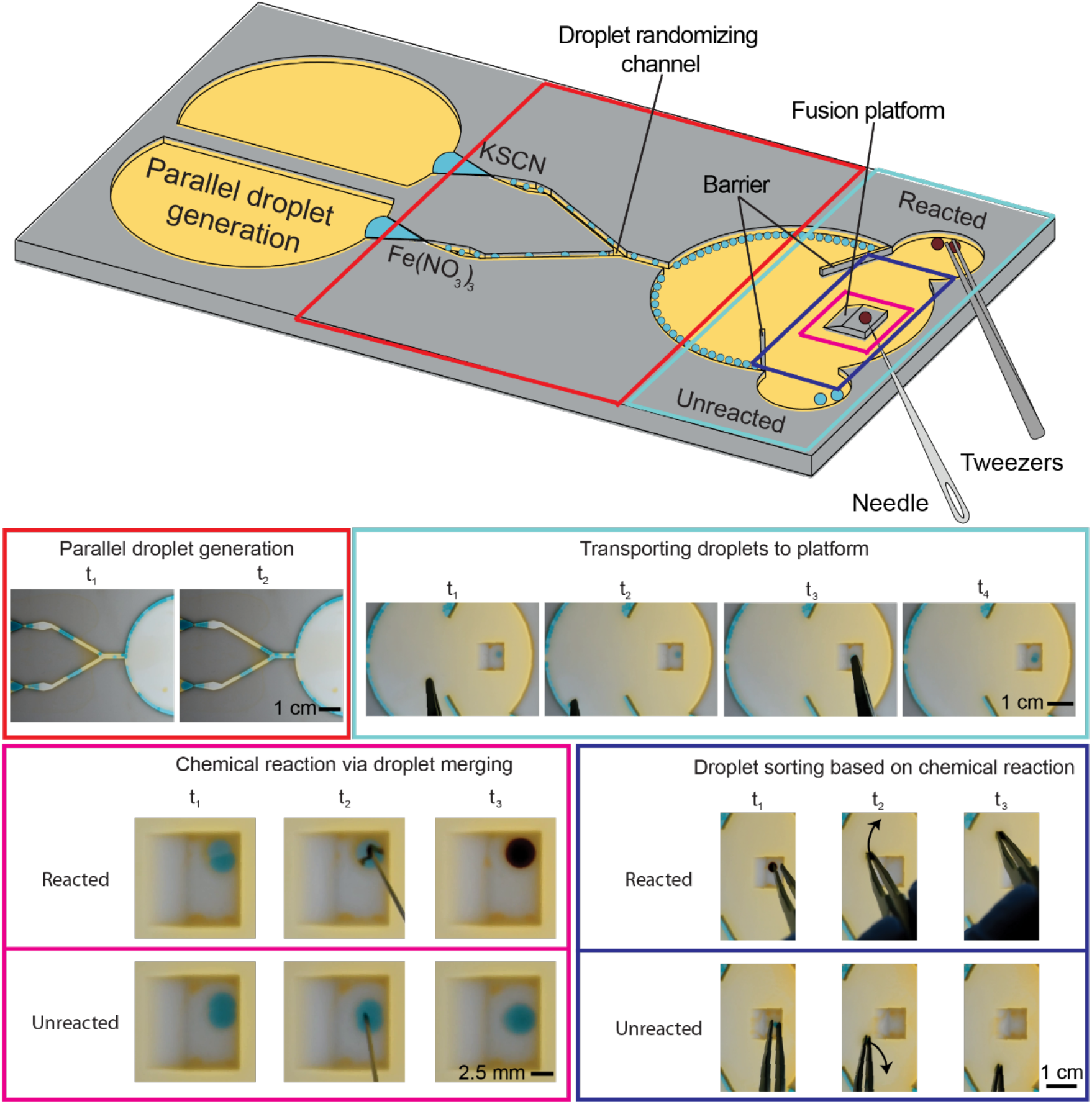
Model chemical reaction using capillary-driven open droplet microfluidics. Droplets containing KSCN and Fe(NO_3_)_3_ are generated in parallel. Two droplets are randomly selected and moved to the fusion platform with tweezers. The droplets are merged with a needle; if a colored complex (Fe(SCN)_3_) forms, the droplet is sorted to the reacted chamber; droplets without Fe(SCN)_3_ are sorted to the unreacted chamber. Video available in Supplementary Video 7.

The purpose of this proof-of-concept workflow is to demonstrate the ability to perform a reaction and sort droplets based on “hits” (in this case droplets containing the reaction product Fe(SCN)_3_). This suggests the potential for conducting droplet sorting tasks in applications like single cell encapsulation, directed evolution of enzymes, and process optimization of synthetic reactions to synthesize small molecules or novel materials.

## Discussion

Overall, our open droplet microfluidic system is the first of its kind that can autonomously generate droplets by utilizing hydrostatic pressure and capillary pressure. Further, our open system enables manipulation of droplets (including patterning, transferring, splitting, merging, and sorting) due to unobstructed access to the droplets along the entire length of the device. This offers opportunities that simply are not possible in closed channel devices, where the ceiling obstructs access. We acknowledge that there are limitations of our system —chiefly, the number of droplets generated (2 droplets/second) in our device is not at the level of high throughput droplet microfluidic systems where droplets are generated at thousands of droplets per second. Thus, we envision applications of our method will capitalize on aspects uniquely enabled by the open channel (such as the ability to manipulate droplets with implements like tweezers or a needle and the ability to add and remove solid objects like beads or pieces of tissue) that are suited for medium-throughput experiments (hundreds to thousands of droplets in total). Furthermore, the ability to parallelize the channel without adding pumps to drive the flow allows for the droplet generation to be scaled up without adding additional equipment for each new droplet generator. Additionally, this method is currently limited to use with PTFE. Although many plastics were tested (i.e., PMMA, polystyrene, polypropylene, etc.) with this channel design, only PTFE was successful in droplet generation with fluorinated oil carrier phases due to its high hydrophobicity. Fortunately, PTFE is a biocompatible material and is suitable for biological applications and it has excellent solvent compatibility for chemical applications.(*46*) Our method opens up numerous areas for future work—both on biological and chemical applications and in further understanding the physics underlying droplet formation and droplet transport via the Cheerios effect. Finally, this design is a first step in exploring the space of power-free open droplet microfluidic systems and provides design rules for similar channel designs.

## Materials and Methods

### Channel design and fabrication

The channels were designed in Solidworks (Dassault Systemes SE) and postprocessed in Fusion 360 (Autodesk, Inc.). Then, they were fabricated out of 3.2 mm thick PTFE (McMaster-Carr Supply Co.) on a Datron NEO mill (Datron Dynamics Inc.). After the channels were milled, they were rinsed with deionized water (DIW), sonicated in 70% ethanol for 30 min, and then rinsed again with DIW. Finally, the channels were dried with compressed air.

The channel height is 1.6 mm, and the width of the narrow constriction ranges between 0.2 mm and 3.0 mm. There is also a platform with a height of 0.2 mm at the narrow constriction and converging-diverging region to allow the aqueous plug to be pinned at the constriction entrance to produce consistent droplet generation. The channel has a square cross section. The converging part of the tapered channel is 45°, and diverging is 20° as these angles were found to be the best at generating monodisperse droplets consistently. The square protrusions that form the channel constriction are 1 mm wide. CAD files and engineering drawings (SI Supplementary Note 4, Figures S3-S6) are included in the SI.

### General reagents

Carrier fluids used were pure HFE 7500 (The 3M Co.), HFE 7500 with 0.002 wt%, 0.02 wt%, 0.2 wt%, and 2 wt% Rhodamine fluorosurfactant (RAN Biotechnologies Inc.). The Rhodamine fluorosurfactant (FS) is a mix of fluorosurfactant 008FS (008-FluoroSurfactant) and Rhodamine-functionalized fluorosurfactant Rhod-FS (FS-Rhodamine) (FS is a mix of 008FS and Rhod-FS with weight ratio of 3:1 respectively). The aqueous phase, DIW, is mixed with blue food dye (McCormick Corp.).

### Image acquisition setup

A Nikon-D5300 DSLR camera attached to a jig with adjustable distance was used to record the experimental results at the top view of the channel and at a frame rate of 30 fps. For the high-speed video in Figure 2b, a Chronos 1.4 high-speed camera (Kron Technologies Inc.) was installed on a stereoscope (United Scope LLC) to record the droplet generation at 2577 fps.

### Droplet generation

A 180 μL DIW plug containing food dye was pipetted on the platform in the channel and then 2 mL of carrier fluid was pipetted into the inlet reservoir of the channel to generate water droplets ranging from 0.52 μL to 34.55 μL (see Figure 1d). Generated droplet volume takes 1 – 2 minutes to stabilize and generate consistent droplet volume.

It was found that droplet generation is initiated faster when the channel is placed at a slight incline of 3°. Placing the channel on an incline did not result in a significant change in the generated droplet volume. Thus, in the proof of concept demonstrating the sorting of reacted products in droplets (Figure 8), the channel was placed at an incline.

### Contact angle and interfacial tension measurements

Contact angles and interfacial tension values reported in Figure 3 were measured on a Kruss Drop Shape Analyzer model DSA025 (Kruss GmbH). Contact angles of the aqueous droplet on the PTFE substrate were measured using the sessile droplet method. When needed, the sessile droplet method is conducted with the sessile droplet submerged in a second liquid to measure the contact angle in a quartz cuvette. In this experiment, the droplet was DIW with blue dye submerged in HFE7500 at various FS concentrations (0 wt%, 0.002 wt%, 0.02 wt%, 0.2 wt%, or 2 wt%) and the substrate was PTFE.

Interfacial tension was measured using the pendant drop method where the fluid body is deposited by a motorized syringe pump until it achieves a pendant shape. From deriving the force balance between the interfacial tension and gravity, the interfacial tension is extracted from the droplet shape. When needed, the pendant drop method is conducted with the pendant drop submerged in a second liquid to measure the interfacial tension in a quartz cuvette. In this experiment, the droplet was DIW with blue dye submerged in HFE7500 at various FS concentrations (0 wt%, 0.002 wt%, 0.02 wt%, 0.2 wt%, or 2 wt%).

### Droplet volume determination

A custom MATLAB script was written to determine the droplet volume. Using image processing, droplets were identified in the experimental videos and the area they encompass was calculated by counting the number of pixels of the droplet. From there, we calculated the effective diameter of the droplets from the area and depending on if droplet diameter was larger than channel height h, the droplets were approximated as either spherical or cylindrical shape. When droplet diameter is less than the channel height, droplets have a spherical-like shape and when droplet diameter is larger than the channel height, droplets have a cylindrical-like shape. To calculate droplet volume:

If droplet diameter is larger than channel height h,

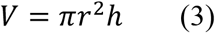

If droplet diameter is smaller than channel height h,

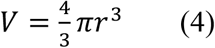

N ≥ 90 droplets were analyzed per data point. The mean (markers) and standard deviation (error bar) of the droplet volume are plotted in Figure 4.

This method of computing droplet volume is an estimation from fitting droplets to simple shapes (sphere and cylinder) and was determined to be satisfactory (See Supplementary Figure S7, *R*^2^ = 0.9799) for this work as the droplets do not spread much. As can be seen in Supplementary Figure S7, the Matlab code computed small droplet volume accurately while larger computed droplet volumes were overestimated. The overestimation is due to the droplets spreading out as Bond number increases making its droplet height shorter which is not accounted for by the code.

### Droplet sorting and merging (Figure 6 & 7)

For transporting a droplet, either a pair of tweezers or a PTFE bead attached to a stylus was used. The stylus was 3D-printed with grey resin (RS-F2-GPGR-04, Formlabs Inc.) and has a ⅛” PTFE ball (McMaster-Carr Supply Co.) adhered to the tip with silicone sealant (Gorilla, Maine Wood Concepts Inc.). The tweezers were a pair of PTFE tweezers with slim, rounded, and smooth tips (McMaster-Carr Supply Co.). For merging droplets, a stainless-steel stitching needle (AEHO crafts) or medical grade No. 22 hypodermic needle was used.

### KSCN and Fe(NO_3_)_3_ aqueous solution preparation (Figure 8)

Green food dye (McCormick Corp.) was added to solutions of 0.1 M of KSCN (Thermo Fisher Scientific Corp.) and 0.05 M of Fe(NO_3_)_3_ (Thermo Fisher Scientific Corp.) until they were approximately the same shade of blue. KSCN is clear while Fe(NO_3_)_3_ is yellow which is why the blue food dye was not added by exact measurements. The workflow of droplet generation, selection, fusion, and sorting (Figure 8) is described in the results section.

## Author contributions

Designed the research: J.K., U.N.L., J.B., E.B., and A.B.T.

Theory discussion and derivation: J.K. and J.B.

Performed experiments: J.K. and U.N.L.

Interpretation of results: J.K., U.N.L., J.B., E.B., and A.B.T.

Writing: J.K., U.N.L., J.B., E.B., and A.B.T.

## Data and materials availability

All data needed to evaluate the paper are presented in the paper and the Supplementary Materials. Additional data may be requested from authors.

