## Supplementary material for "Interfacial tension driven open droplet microfluidics": Biorxiv_supplementary_materials_aliquot.pdf

**This PDF file includes:**

**Supplementary figures and tables**

Supplementary Figure S1. Regions of the device
Supplementary Figure S2. Physical parameters for aqueous plug extrusion
Supplementary Table S1. Table of parameters of Figure 3a regime map
Supplementary Table S2. Table of parameters of Figure 4 generated droplet volume
Supplementary Figure S3. Engineering drawing of droplet generation channel
Supplementary Figure S4. Engineering drawing of sorting channel. (Note: CAD file is also included in the uploaded SI files.)
Supplementary Figure S5. Engineering drawing of UW channel. (Note: CAD file is also included in the uploaded SI files.)
Supplementary Figure S6. Engineering drawing of fusion channel. (Note: CAD file is also included in the uploaded SI files.)
Supplementary Figure S7.

**Other Supplementary Materials for this manuscript include the following:**
**CAD Files**

CAD File of droplet generation channel
CAD File of sorting channel
CAD File of UW channel
CAD File of fusion channel

**Videos**

Supplementary Video 1 Droplet generation
Supplementary Video 2 Droplet patterning around UW shape
Supplementary Video 3 Droplet transferring
Supplementary Video 4 Droplet merging

Supplementary Video 5 Droplet sorting
Supplementary Video 6 Droplet splitting
Supplementary Video 7 Fusion channel

### Supplementary Text

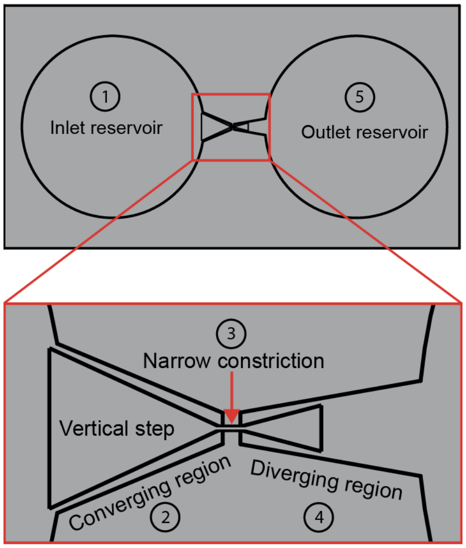


**Supplementary Figure S1.** Main regions of the device. (1) Inlet reservoir which leads to the (2) converging region, followed by the (3) narrow constriction, (4) diverging region, and (5) outlet reservoir.

**Supplementary Note 1: Theoretical derivation of condition for aqueous plug extrusion or droplet generation in our channel**

In this section, a theoretical model is derived to determine the condition required for aqueous plug extrusion or droplet generation in our channel. Let us consider the schematization of **Supplementary Figure S2** where an aqueous plug is placed in a constriction and both sides of the plug are filled with an immiscible carrier fluid. The converging region holds the aqueous plug where the anterior end of the plug meets the narrow constriction. The pressure on the anterior and posterior sides of the plug are different. The model is based on the fact that the pressure difference between the posterior and anterior interfaces of the plug causes the plug extrusion or droplet generation. This pressure difference is due to the hydrostatic pressure exerted by the carrier in the reservoir on the aqueous plug. In our system, the interfacial tension between carrier fluid and aqueous plug (HFE7500/deionized water) is very low. Therefore, the carrier fluid (HFE7500) does not flow over the aqueous plug. This configuration is called shift mode as defined in Lee *et al.*[1] Furthermore, because the aqueous plug does not wet the PTFE channel walls, there is a small leakage of carrier fluid along the inner corners of the converging region.[2] It is assumed that this leakage of carrier fluid imposes the same interfacial tension on both corners (left and right) of the aqueous plug. Finally, a simplifying assumption has been made: The vertical curvature radii on both sides of the plug cancel out.

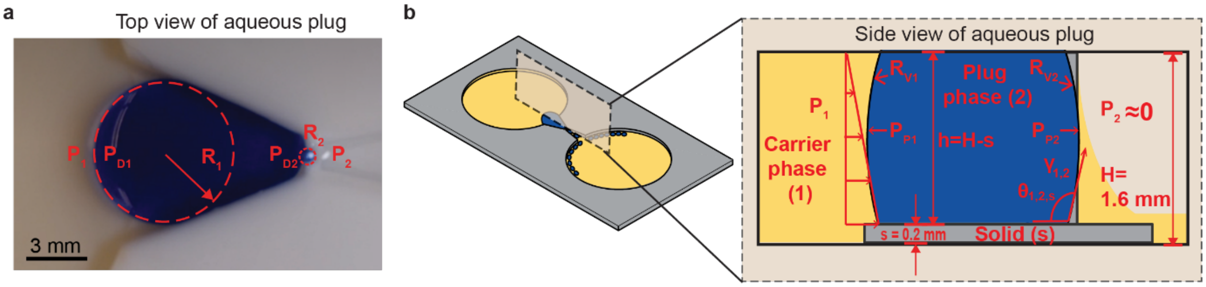


**Supplementary Figure S2**. **a** Top and **b** side view of aqueous plug placed in the converging region prior to aqueous plug extrusion.

Let us write the Laplace pressure for each end of the aqueous plug[2-4]

For the posterior interface,

$$P_{P1} - P_1 = \gamma \left(\frac{1}{R_1} + \frac{1}{R_{v1}}\right) \quad (1)$$

Similarly for the anterior interface,

$$P_{P2} - P_2 = \gamma \left(\frac{1}{R_2} + \frac{1}{R_{v2}}\right) \quad (2)$$

Where $P_{Pi}$ is the local pressure in the aqueous plug at the interface, $P_i$ is the carrier fluid pressure at the external aqueous plug interface, $\gamma$ is the interfacial tension between carrier fluid and aqueous plug, $R_i$ is the curvature radius in the horizontal plane (**Supplementary Figure S2a**), and $R_{vi}$ is the curvature radius in the vertical plane (**Supplementary Figure S2b**). When the subscript of parameters, $i = 1$, it denotes that the parameter is referring to parameters at the posterior side of the aqueous plug and when $i = 2$, it denotes that the parameter is referring to parameters at the anterior side of the aqueous plug.

Subtracting Eq. (2) from Eq. (1) yields

$$P_{P1} - P_{P2} = (P_1 - P_2) + \gamma \left(\frac{1}{R_1} - \frac{1}{R_2} + \frac{1}{R_{v1}} - \frac{1}{R_{v2}}\right) \quad (3)$$

We make the assumption that $R_{v1} \approx R_{v2}$ because the channel height is uniform throughout the channel and we are left with

$$P_{P1} - P_{P2} \cong (P_1 - P_2) + \gamma \left(\frac{1}{R_1} - \frac{1}{R_2}\right) \quad (4)$$

The pressure in the inlet reservoir $P_1$ (see **Supplementary Figure S2a**) is the hydrostatic pressure ($P_1 = \rho g h$) and that in the outlet, nearly empty reservoir is determined to be $P_2 \approx 0$. Then relation (4) becomes

$$P_{P1} - P_{P2} \cong \rho g h + \gamma \left(\frac{1}{R_1} - \frac{1}{R_2}\right) \quad (5)$$

where $\rho$ is the density of the carrier fluid, $g$ is the gravitational acceleration, and $h$ is the carrier fluid height (note that for our device, aqueous plug height is also equal to carrier fluid height).

The condition for the aqueous plug to advance into the constriction is $P_{P1} > P_{P2}$, or

$$\rho g h + \gamma \left(\frac{1}{R_1} - \frac{1}{R_2}\right) > 0 \quad (6)$$

At the beginning of the plug advancement (depicted in Figure 2), the curvature radius $R_1$ is very large when compared to curvature radius $R_2$ ($R_1 \gg R_2$). This means $1/R_1$ can be neglected when subtracted by $1/R_2$ ($1/R_1 \ll 1/R_2$ $so$ $1/R_1 - 1/R_2 \approx -1/R_2$).

Then the condition for plug extrusion or droplet generation is

$$\rho g h > \frac{\gamma}{R_2} \quad (7)$$

When the anterior section of the aqueous plug interface advances through the constriction (see **Supplementary Figure S2a**), the initial curvature radius $R_2$ is approximately[2]

$$R_2 \approx \frac{w}{2|cos\theta|} \quad (8)$$

where $w$ is the constriction width and $\theta$ is the contact angle between channel wall, aqueous plug, and carrier fluid.

The condition for plug extrusion or droplet generation is then

$$\rho g h > \frac{2\gamma\ |cos\theta|}{w} \qquad (9)$$

Relation Eq. (9) shows that the constriction width *w* must not be too small when designing the device ($w\ > \frac{2\gamma\ |cos\theta|}{\rho g h}$).

**Supplementary Note 2: Use of units when computing $\frac{2\gamma\ |cos\theta|}{\rho g h}$ in Figure 3**

When calculating $\frac{2\gamma\ |cos\theta|}{\rho g h}$, we use SI units for all parameters during calculation; thus, the final value will be the SI unit for length, meters. In Figure 3 since we used millimeters on the x-axis, we convert $\frac{2\gamma\ |cos\theta|}{\rho g h}$ to millimeters (1 mm = 0.001 m).

Here, we show the rearrangement and simplification of the units of $w\ > \frac{2\gamma\ |cos\theta|}{\rho g h}$ and show that the final unit is meter.

The unit of $\gamma$ is of N/m, $\theta$ is unitless, $\rho$ is kg/m^3, g is m/s^2, and h is m. Unit of N is kg*m/s^2

$\frac{2\gamma\ |cos\theta|}{\rho g h}$ would then have units:

$$\frac{\frac{N}{m}}{\left(\frac{kg}{m^3}\right)*\left(\frac{m}{s^2}\right)*(m)} = \frac{N*m^3*s^2}{kg*m^3} = \frac{N*s^2}{kg} = \frac{kg*m*s^2}{kg*s^2} = m$$

From there, convert meters to millimeters to obtain the same unit as Figure 3a.

**Supplementary Note 3: Table of parameters**

**Supplementary Table S1.** Table of parameters of Figure 3a regime map.

| Constriction width (mm) | Interfacial tension (mN/m) | Density (kg/m^3) | Carrier fluid height (mm) | Contact angle (degree) | Plug Extrusion (Yes/No) | Droplet generation (Yes/No) |
|---|---|---|---|---|---|---|
| 0.20 | 3.9 | 1614 | 1.4 | 125.2 | Yes | Yes |
| 0.40 | 3.9 | 1614 | 1.4 | 125.2 | Yes | Yes |
| 0.60 | 3.9 | 1614 | 1.4 | 125.2 | Yes | Yes |
| 3.00 | 3.9 | 1614 | 1.4 | 125.2 | Yes | No |
| 1.00 | 3.9 | 1614 | 1.4 | 125.2 | Yes | No |
| 2.00 | 3.9 | 1614 | 1.4 | 125.2 | Yes | No |
| 0.20 | 10.3 | 1614 | 1.4 | 132.4 | No | Yes |
| 0.40 | 10.3 | 1614 | 1.4 | 132.4 | Yes | Yes |
| 0.60 | 10.3 | 1614 | 1.4 | 132.4 | Yes | Yes |
| 3.00 | 10.3 | 1614 | 1.4 | 132.4 | Yes | No |
| 0.25 | 14.8 | 1614 | 1.4 | 135.2 | No | No |
| 0.40 | 14.8 | 1614 | 1.4 | 135.2 | No | No |
| 1.00 | 14.8 | 1614 | 1.4 | 135.2 | No | No |
| 2.00 | 14.8 | 1614 | 1.4 | 135.2 | Yes | No |
| 3.00 | 14.8 | 1614 | 1.4 | 135.2 | Yes | No |
| 1.00 | 35.0 | 1614 | 1.4 | 140.1 | No | No |
| 2.00 | 35.0 | 1614 | 1.4 | 140.1 | No | No |
| 3.00 | 35.0 | 1614 | 1.4 | 140.1 | No | No |
| 0.25 | 35.0 | 1614 | 1.4 | 140.1 | No | No |
| 0.40 | 35.0 | 1614 | 1.4 | 140.1 | No | No |
| 0.20 | 47.9 | 1614 | 1.4 | 147.2 | No | No |
| 0.40 | 47.9 | 1614 | 1.4 | 147.2 | No | No |
| 0.60 | 47.9 | 1614 | 1.4 | 147.2 | No | No |
| 3.00 | 47.9 | 1614 | 1.4 | 147.2 | No | No |
| 1.00 | 47.9 | 1614 | 1.4 | 147.2 | No | No |
| 2.00 | 47.9 | 1614 | 1.4 | 147.2 | No | No |

**Supplementary Table S2.** Table of parameters of Figure 4 generated droplet volume

| Constriction width (mm) | Density (kg/m^3) | Carrier fluid height (mm) | Interfacial tension (mN/m) | Contact angle (degree) | Generated droplet volume (μL) | |
|---|---|---|---|---|---|---|
| | | | | | Average | Standard deviation |
| 0.20 | 1614 | 1.4 | 3.9 | 125.2 | 0.52 | 0.025 |
| 0.40 | 1614 | 1.4 | 3.9 | 125.2 | 2.49 | 0.175 |
| 0.60 | 1614 | 1.4 | 3.9 | 125.2 | 7.75 | 0.465 |
| 1.00 | 1614 | 1.4 | 3.9 | 125.2 | 19.22 | 1.344 |
| 0.40 | 1614 | 1.4 | 10.3 | 132.4 | 7.23 | 0.504 |
| 0.60 | 1614 | 1.4 | 10.3 | 132.4 | 15.09 | 0.906 |
| 1.00 | 1614 | 1.4 | 10.3 | 132.4 | 34.55 | 2.422 |

### Supplementary Note 4: Engineering drawing of open microfluidic channels

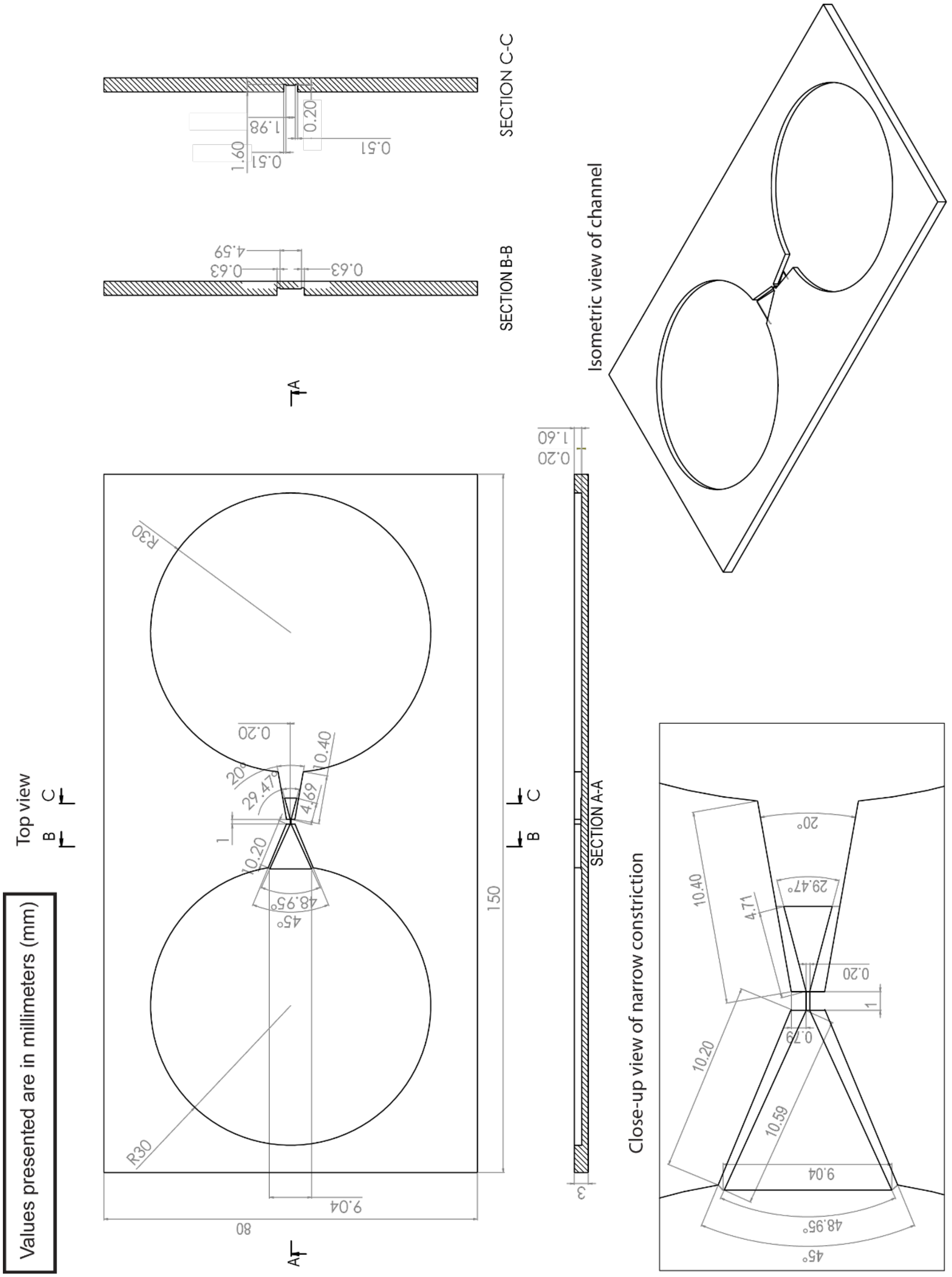


**Supplementary Figure S3.** Engineering drawing of droplet generation channel. (Note: CAD file is also included in the uploaded SI files.)

Values presented are in millimeters (mm)

Detailed measurements of converging-diverging region and constriction can be found in Supplementary figure S3

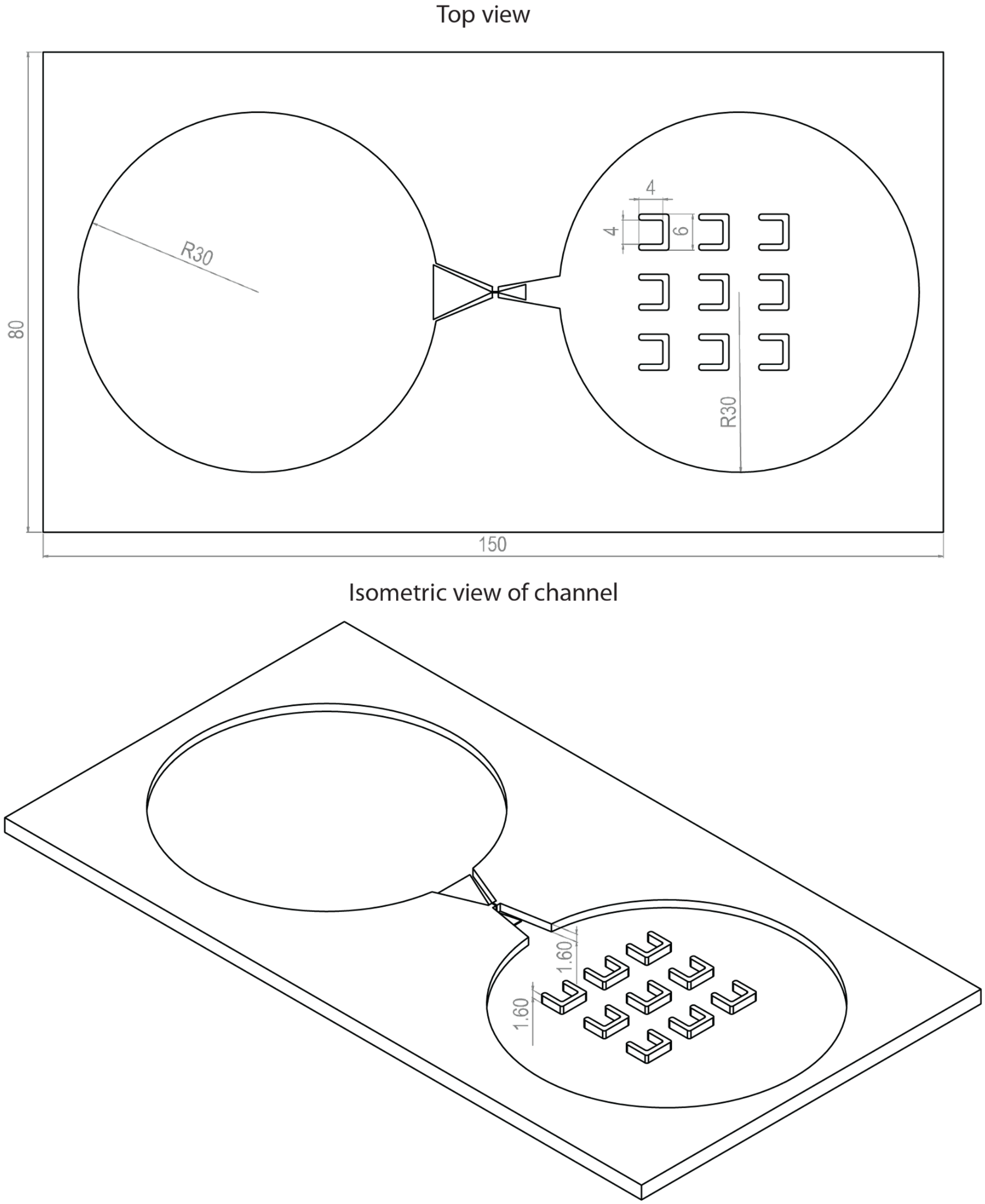


**Supplementary Figure S4.** Engineering drawing of sorting channel. (Note: CAD file is also included in the uploaded SI files.)

Values presented are in millimeters (mm)

Detailed measurements of converging-diverging region and constriction can be found in Supplementary figure S3

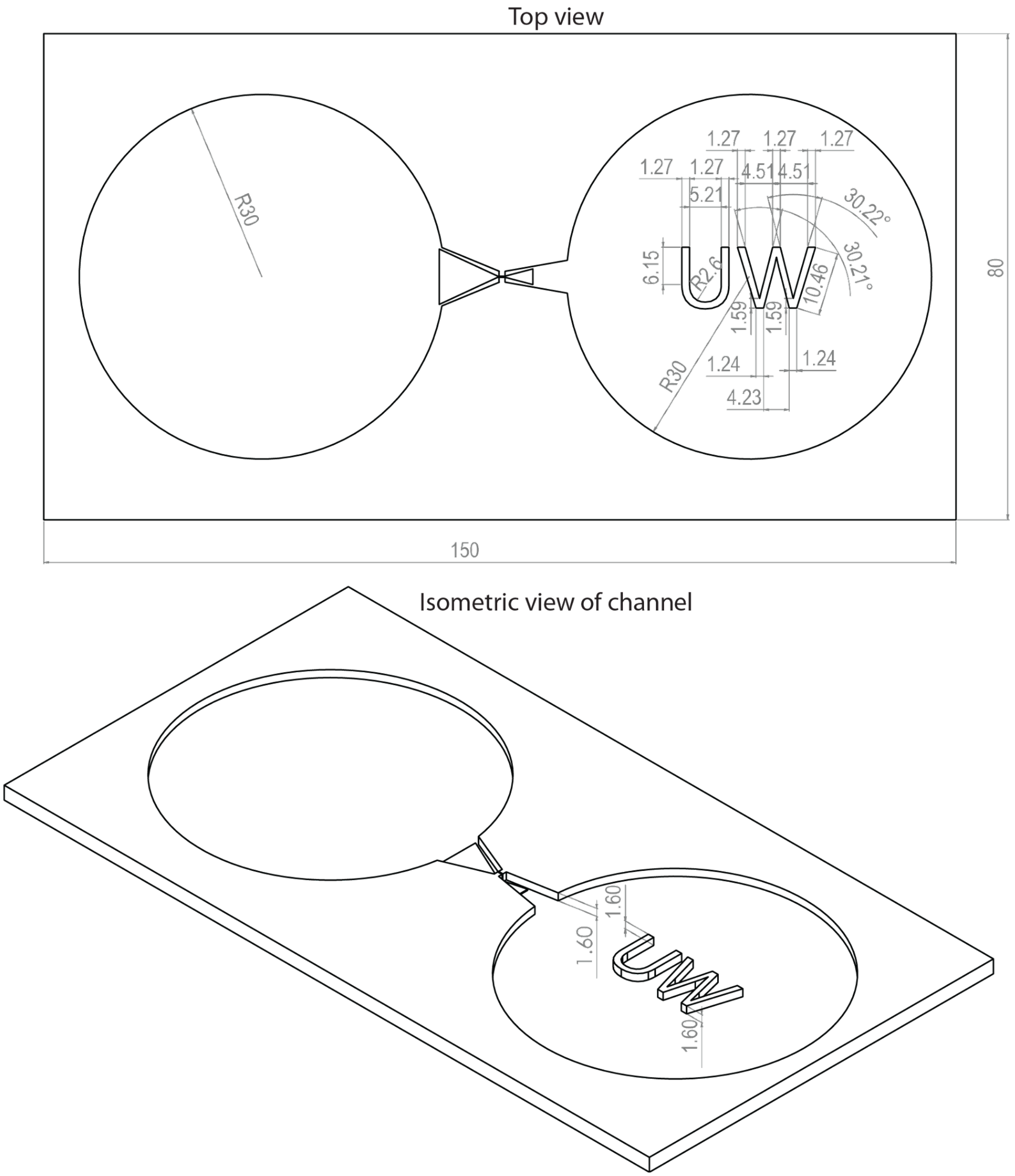


**Supplementary Figure S5.** Engineering drawing of UW channel. (Note: CAD file is also included in the uploaded SI files.)

Values presented are in millimeters (mm)

Detailed measurements of converging-diverging region and constriction can be found in Supplementary figure S3

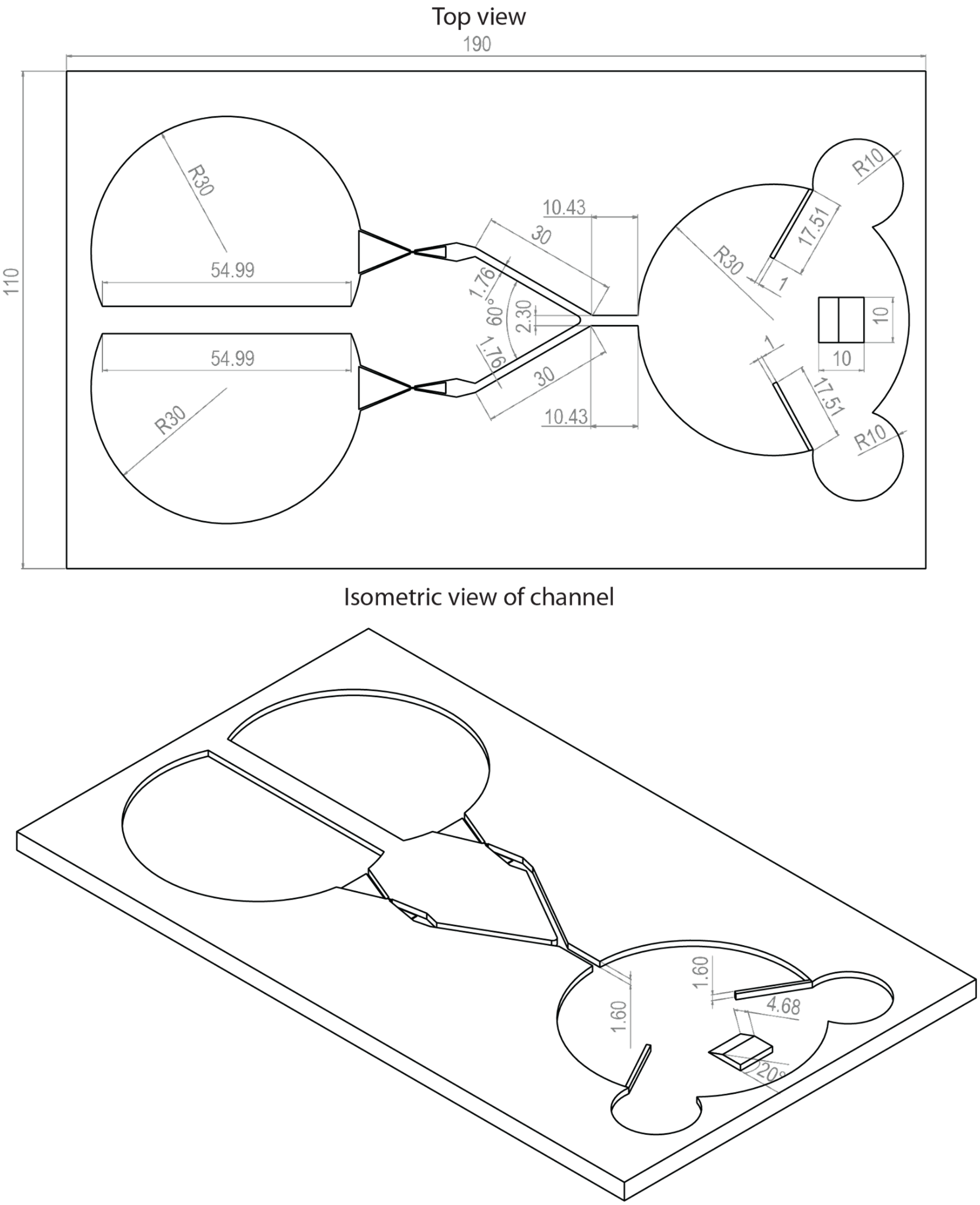


**Supplementary Figure S6.** Engineering drawing of fusion channel. (Note: CAD file is also included in the uploaded SI files.)

**Supplementary Note 5: Accuracy of Matlab code during computation of droplet volume.**

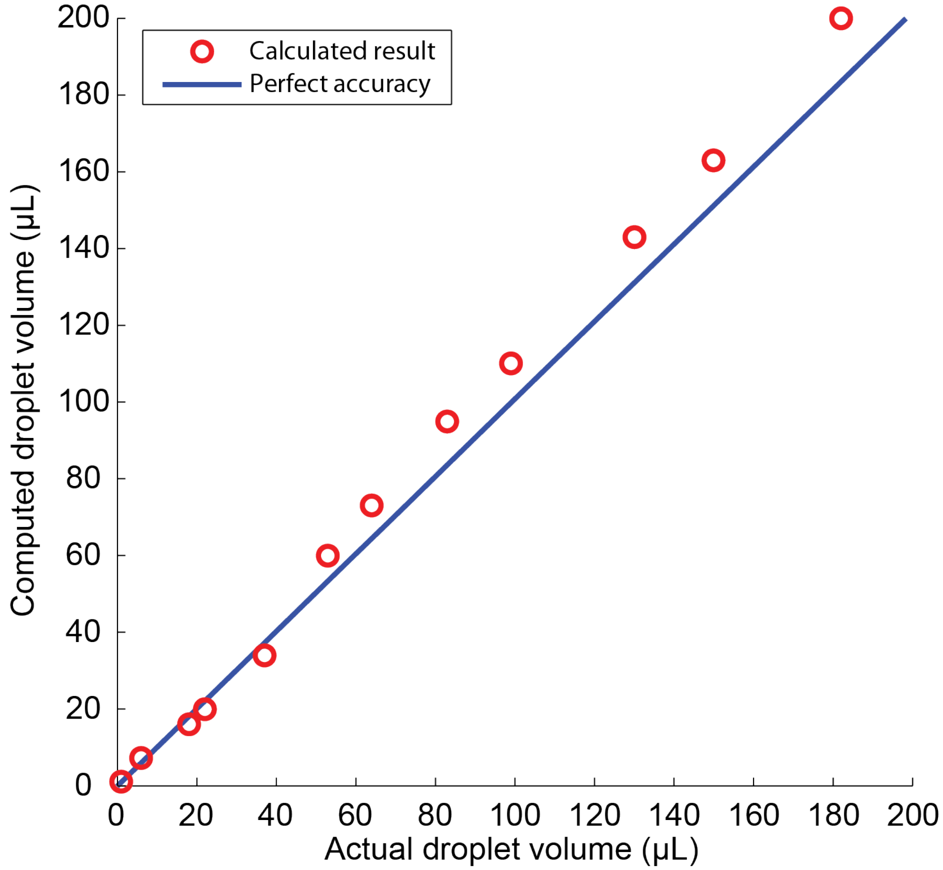


**Supplementary Figure S7.** Plot that shows the accuracy of the custom Matlab code in determining droplet volume when compared to actual droplet volume. Red circles represent droplet volume computed by Matlab code and blue solid line represents the curve when the Matlab code would achieve a perfect accuracy (100%). When compared to the "perfect accuracy" reference curve, the data points has coefficient of determination, $R^2 = 0.9799$.
